# A seven-year record of fluctuating core body temperatures of nesting leatherback and hawksbill sea turtles

**DOI:** 10.1101/2023.07.28.551020

**Authors:** Malcolm W. Kennedy

## Abstract

Sea turtles experience changes in water temperatures during migrations and seasonal movements that will influence their body temperatures. Nothing is known of how sea turtles’ core body temperatures vary from season to season at nesting sites. Over seven consecutive seasons we measured the surface temperatures of freshly laid eggs as proxies of core temperatures of sea turtles using non-contact infrared thermometers. We measured egg temperatures of two species that have contrasting lifestyles - leatherbacks, the largest, which are adapted to migrate between tropical breeding sites to cold temperate waters, and the smaller hawksbills that are confined to the tropics and sub-tropics. We found considerable year-to-year variations in temperatures in both species (year means 30.4 °C to 31.5 °C in leatherbacks), hawksbills the more so (28.1 °C to 30.3 °C). These differences will likely be modified by both natural seasonal variations and anthropogenic changes in global ocean temperatures and resulting changes in currents and water temperatures local to nesting beaches. These previously unrecognised diversities in body temperatures of nesting turtles are pertinent to predicting environmental tolerances, reproductive success, and nest site selection by sea turtles, and could contribute to predicting which rookeries may remain viable or not during future ocean warming.

## 1.1 Introduction

Sea turtles range across the oceans for feeding and breeding, facing a broad range of environmental challenges. Those that encounter the greatest temperature ranges are adult leatherback turtles, *Dermochelys coriacea*, which travel from the tropics where they mate and nest, to cold temperate waters where many feed between nesting [1-5]. Leatherbacks survive and forage in sea temperatures that would cold-stun other species [5-8]. They achieve this through extensive fat storage for thermal insulation, countercurrent vascular systems in their limbs and the muscles that operate them, and their sheer size (“gigantism”) confers a reduced surface area to volume ratio through which to lose heat [9-13].

Cumulatively, these heat conservation features allow them to retain core body temperatures consistently above ambient sea levels [6, 14]. All other sea turtle species, such as hawksbills, are mainly confined to tropical and subtropical waters, and do not encounter similar extremes.

Rising anthropogenic global temperatures are leading to increasing concerns about how and where sea turtles may successfully nest, or indeed survive, in the seas around, and migration routes to, their rookeries [15-19]. There are already predictions that oceans may warm to a degree that may be intolerable to sea turtles, the well-insulated leatherbacks being therefore at particular risk [14, 20, 21]. Moreover, rising sand temperatures on nest beaches are increasingly thought to be to the detriment of egg survival [22-27], and, through temperature-dependent sex determination during development in chelonians, deemed to already be affecting sex ratios of hatchlings such that an excess of females appears [14, 25, 28].

Increased temperatures of beaches may or may not be paralleled by increases in adjacent seas - changes in ocean temperatures may de-stabilise sea currents and eddies that may thereby change the temperatures, flow rates, and directions of waters local to nesting beaches, in addition to sea level rise [29-33]. Changes in ocean currents caused by global warming may therefore bring warmer or cooler currents to increasingly hot beaches. Were sea turtles to choose a nesting beach partly on the temperatures of the waters they traverse during migration, or in which they congregate before heading to shore, then disparities may arise between beaches where turtles choose to nest and where offspring number and sex ratios are optimum.

The degree to which sea turtles will change, or already are changing, their nest sites and migration routes in response to global warming is receiving attention [14, 28, 34, 35]. Predicting which beaches and the sea approaches to them are to become reproductively unstable, or simply avoided by turtles, will require an understanding of how they respond to increasingly variable ocean conditions. As global warming proceeds, nesting beaches may also oscillate between being non-viable one year to viable the next. Work at several sites has been following seasonal and annual fluctuations in beach temperatures, and temperatures within nests, including studying the gradient of temperatures vertically through nests, of several species of sea turtle [28, 36-38]. Meanwhile, high resolution satellite-based data on sea surface temperatures are informative of changing conditions, although this is complicated by differences between surface and bulk sub-surface water [29, 30, 39-43]. Moreover, given the complexity of water movements in inshore waters, it will likely be difficult to gauge the temperatures of waters in which turtles sojourn before landing to nest, and how they vary through a season or from year to year.

One way of following the offshore temperature regimes to which sea turtles are exposed, and how they are responding to temperature changes, is to measure the core body temperatures of nesting females soon after they emerge from the sea. These have been measured using potentially harmful invasive techniques (summarised in ref. [44]), but more recently by measuring freshly laid eggs [44]. This assumes that eggs in the oviduct thermally equilibrate to the internal tissues of a turtle, such that measuring the temperatures of eggs within a second or two of laying should act as a proxy for a female’s core body temperature. Moreover, this can be done non-destructively by using infrared-based thermometers, which allows measurement of large numbers of eggs of many females in order to obtain reliable mean values for each individual easily in the field.

Using egg surface temperatures as proxy, we carried out a large scale, seven-year survey of core body temperatures of two species of sea turtle with contrasting migratory propensities and body sizes - leatherback and the smaller hawksbill turtles. Hawksbills do not possess the overt heat-retentive anatomical and physiological features of leatherbacks, so should have body temperatures closer to surrounding water and are taken to be ectotherms [14, 45] and be more likely to fluctuate with it. We found a considerable diversity in body temperatures in both species, most noticeably in hawksbills, and distinct year-to-year variability in both species that were more of note in hawksbills.

## 2. Methods

### 2.1 Study sites

Observations of nesting leatherback sea turtles (*Dermochelys coriacea*) were made on Fishing Pond beach on the east coast of Trinidad (approximately 10.60° N, 61.02° W), and hawksbill sea turtles (*Eretmochelys imbricata*) on Campbleton and Hermitage bays in the northeast of Tobago (approximately 11.31° N, 60.57° W). The sites are approximately 95 km apart. Measurements of egg temperatures were made during early June to early August 2013 to 2019. The measurements began approximately at the peak of nesting activity of each species in Trinidad.

### 2.2 Measurement of core body temperatures

The surface temperature of freshly laid eggs was used as a non-contact proxy of core body temperature as previously described [44]. Measurements were taken using Fluke 62 Max Plus hand held dual laser spot-guided infrared thermometers (spectral range 8–14 μm, accuracy ±1 °C or 1%, thermal sensitivity/resolution 0.1 °C) set to an emissivity of 0.98, which is a value taken as typical for emissivity of biological tissues (McCafferty et al., 2013; Rowe et al., 2013; Mortola, 2013). Infrared thermometers were calibrated against a thermocouple and a mercury thermometer of 0.1 °C sensitivity. For leatherbacks, sand was removed from the edges of the nest hole to gain access to the eggs. For hawksbills, sand was removed from the rear of the nest cavity to the same end. In both cases this could be done with minimal or usually no contact with the animal. Measurements were taken at about 20 cm from the eggs surface, and only on eggs that were clearly observed to be freshly laid, and had no sand attached to the side which was being measured. Measurements were taken within one or two seconds of emergence of an egg to avoid any significant cooling by evaporation. The egg temperatures used for each turtle sampled are the average of between 5 and about 30 eggs measured per turtle, and there were essentially no differences in the temperatures recorded during an individual animal’s nesting. The values for 2013 and 2014 are the same as those we reported previously [44].

### 2.3 Sea surface temperatures

Daily sea surface temperatures were obtained from satellite-based data (https://oceanwatch.pifsc.noaa.gov/erddap/griddap/CRW_sst_v3_1.html), using the latitude and longitude settings of 10.60559; 298.97712 for the Trinidad site of leatherback turtle nesting (Fishing Pond beach), and 11.31774; 299.43274 for the Tobago hawksbill site (Hermitage and Campbleton beaches). These coordinates were set for approximately 200 m offshore - where the female turtles tarry before coming to shore to nest is not known in either case, hence this approximation.

### 2.4 Statistical procedures

Data were analysed and plotted using Microcal ORIGIN 2022 and MS Excel softwares. Analysis of variance analyses (ANOVA) were carried out using the Tukey procedure. Data smoothing used the default smoothing procedure in ORIGINlab software set for 50 points.

## 3. Results

### 3.1 Body temperatures of leatherback and hawksbill turtles

271 nesting events by leatherback and 312 by hawksbill turtles were recorded over the seven nesting seasons 2013 to 2019. Pooling the data from all seven years yields temperature frequency distributions for each species as given in figure 1 (see also figure S1). Mean core body temperatures for leatherbacks and hawksbills were 30.9°C and 29.2 °C, respectively, with the medians and modes approximately 1.7 °C apart, and the variance for hawksbills being approximately double that of leatherbacks (see statistical summary in figure 1). These temperature values are similar to those reported by others for leatherbacks when in their nesting areas, using both invasive and non-invasive methods (summarised in [44]).

**Figure 1.**
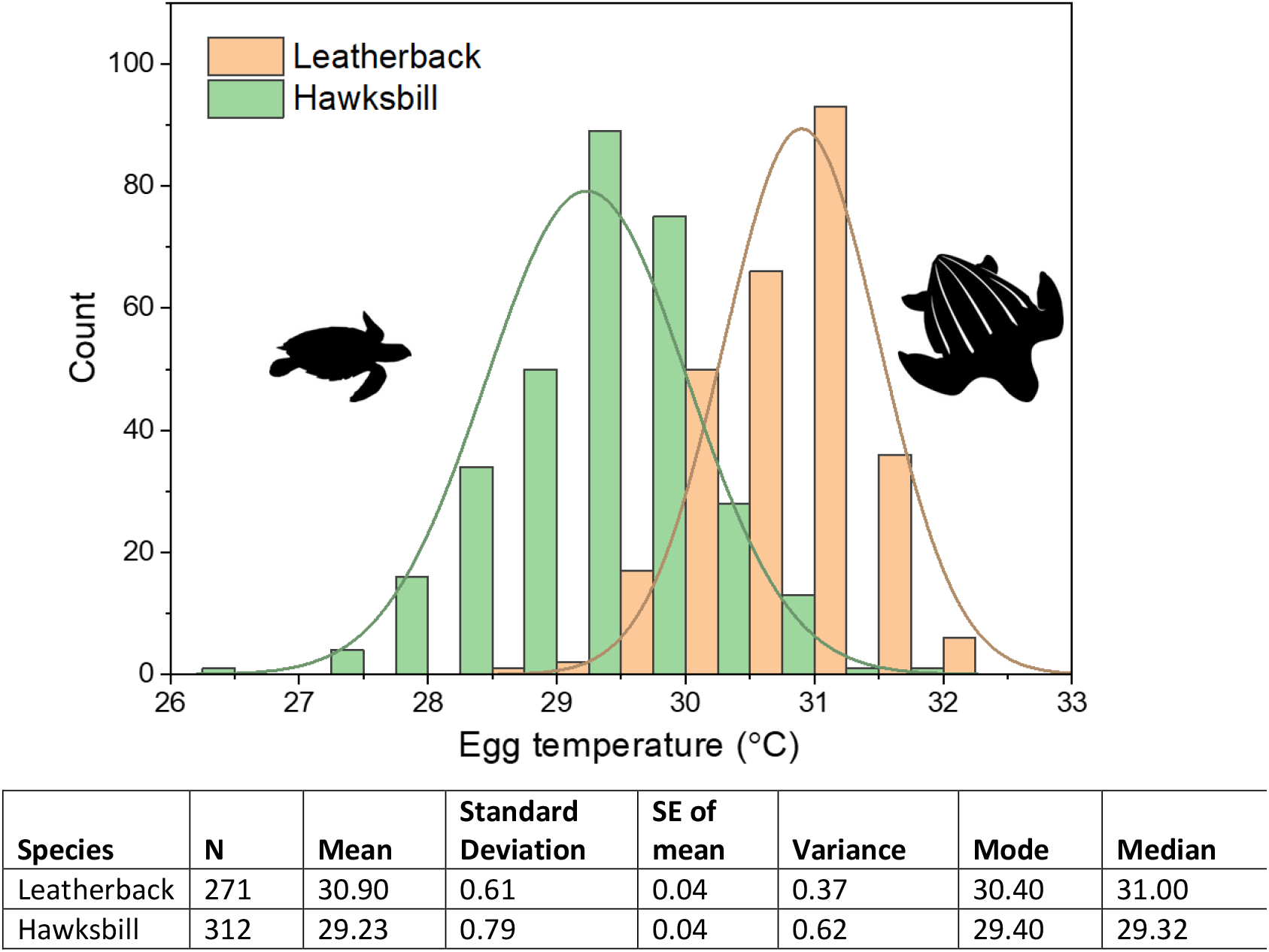
Core body temperature differences between leatherback and hawksbill turtles. Cumulative egg temperature measurements of the two species accumulated over the seven field seasons 2013 – 2019 inclusive. See also figure S1. Data were obtained from 271 nesting instances of leatherback turtles and 312 hawksbills, and a Gaussian distribution is fitted to them separately. Descriptive statistics for the data for each species are given below the figure. Note that the variance for leatherbacks is almost half that for hawksbills. Turtle silhouettes from http://phylopic.org/.

The higher values for leatherbacks over hawksbills are presumably due to their size and anatomical adaptations the former have for metabolic heat retention (see above).

### 3.2 Year to year variation in sea turtle body temperatures

Inspection of the data for the separate nesting season years indicates distinct variation from year to year in both species but is greater for hawksbills (figure 2). Within a given season there were relatively small changes in both leatherback and hawksbill core temperatures as the season progressed, although gradual rises or falls were observed (figure S2). Important here, however, were statistically significant increases or decreases, or none, in core temperatures from year to year for both species (Table S1). Year-to-year fluctuations were greater in hawksbills than leatherbacks (Table S1), potentially due to a combination of intrinsic differences in the species in body temperature regulation or differences in the waters they occupied before coming ashore. Figure 2 also shows that the overall pattern of mean core temperature increases or decreases from year-to-year were similar between the two species, albeit on different scales, dampened in leatherbacks. When the yearly values for each species are plotted against one another there is a good correlation between the two (figure 3A). This is presumably indicative of changing sea temperatures affecting both species, with leatherbacks seemingly more resistant to such fluctuations. However, it could also be that, despite the displacement of the two populations being only 95 km, and both islands being fully exposed to the North Equatorial Current flowing from the east, the waters to which they are exposed may fluctuate locally to different degrees. However, the sea temperatures at the coordinates we chose correlated closely, as indicated by comparing average sea temperatures at each site during the periods of between June 15^th^ and July 15^th^ in each year (approximately covering the central period of the field measurements; figure 3B), or in the overall pattern of fluctuations in daily sea surface temperature recordings (figures 4 and S4).

**Figure 2.**
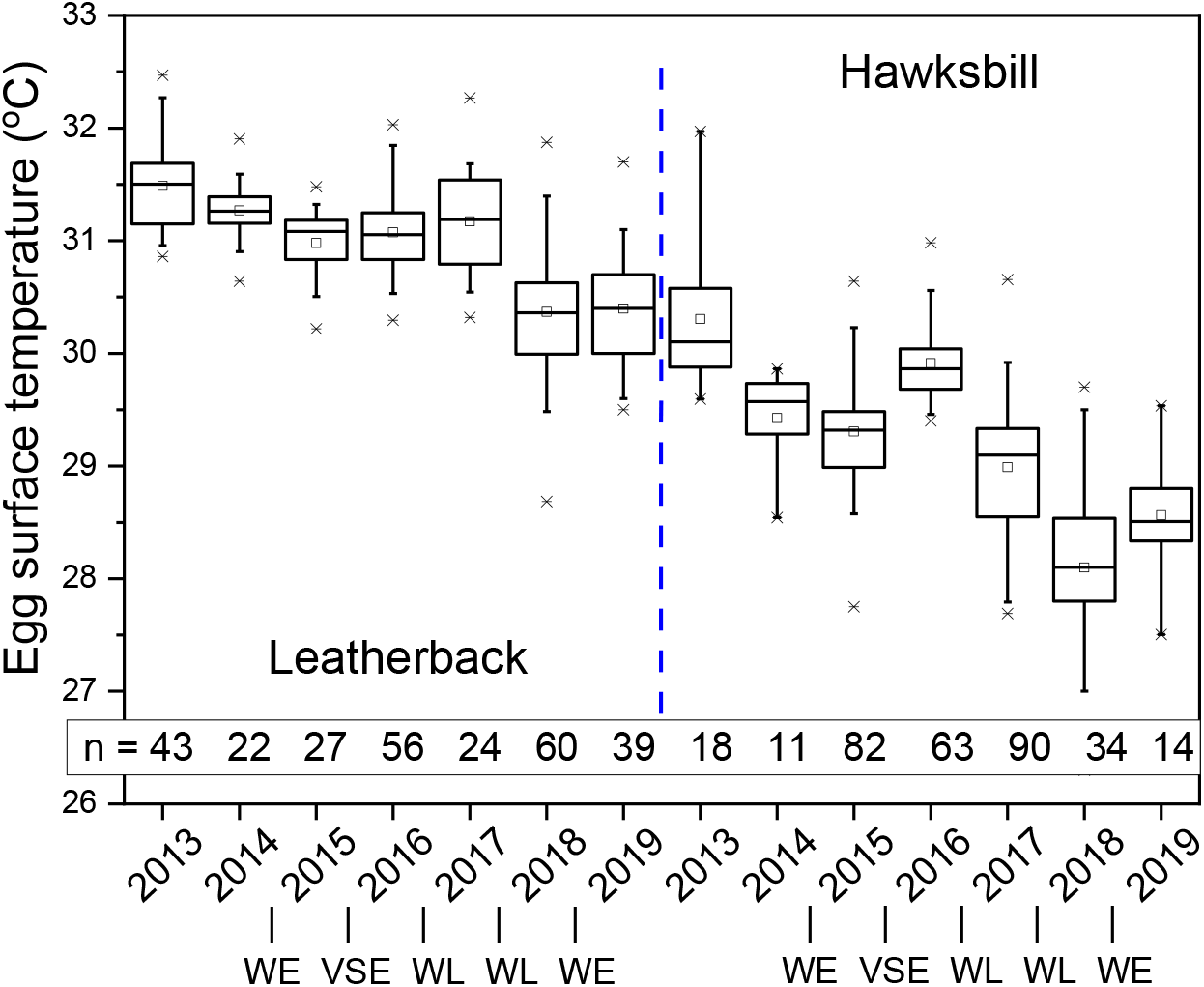
Differences in core body temperatures of leatherback and hawksbill turtles over seven nesting seasons. Boxes represent 25 and 75 % interquartile ranges; horizontal lines through boxes indicate medians; squares indicate the means; whiskers show 5^th^ and 95^th^ percentiles; and crosses indicate the maximum and minimum values. The number of turtle nestings observed in each year are as indicated (n). WE = Weak El Niño; VSE = Very Strong El Niño, WL = Weak La Niña; other years were classified as neutral (https://ggweather.com/enso/oni.htm).

**Figure 3.**
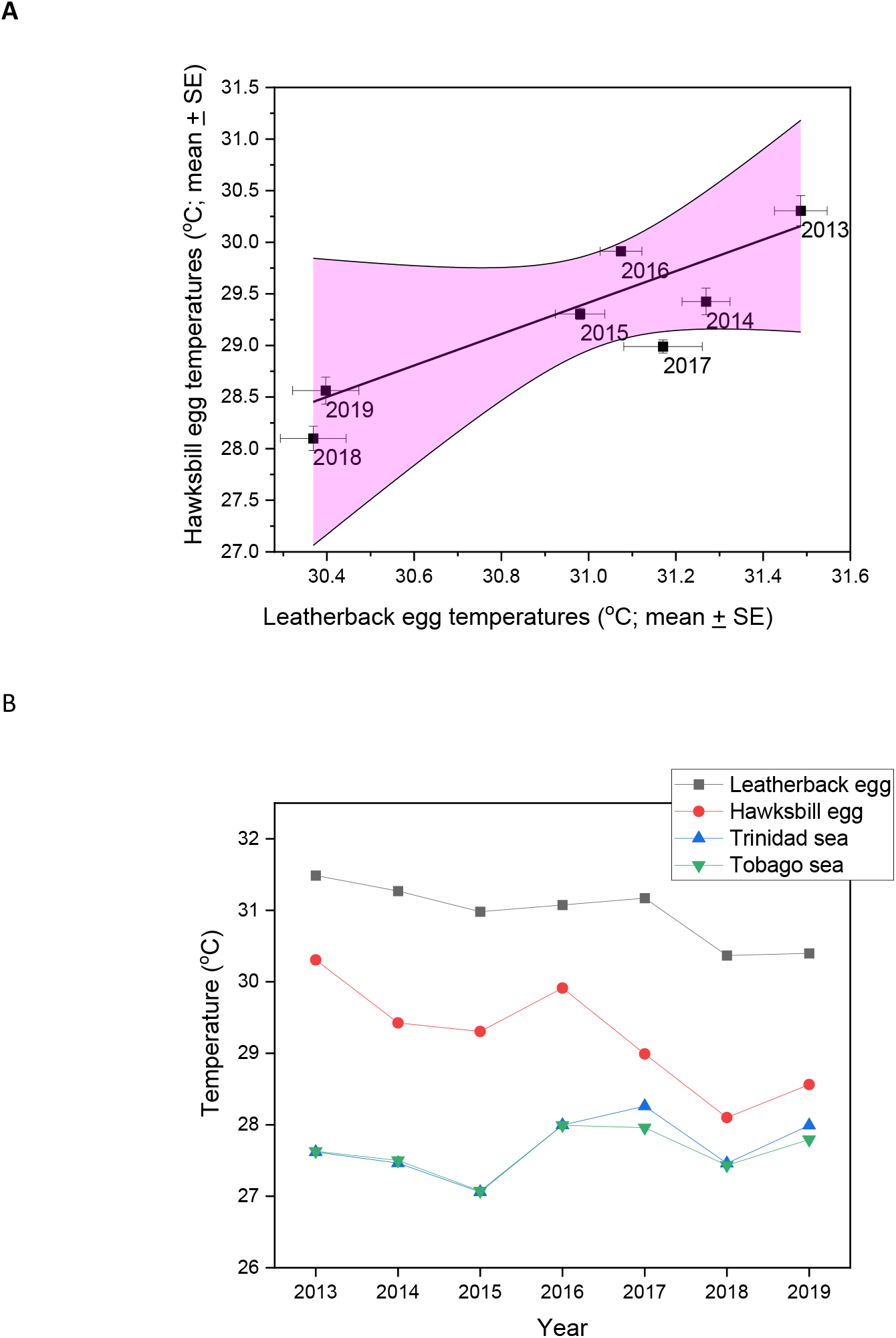
Correlation between female leatherback and hawksbill turtle egg temperatures at the two nesting areas. (A) Average temperatures for all the eggs of each species measured for each year ± standard errors. The linear line fit values are equation y = 1.55x - 18.77; R² = 0.76 and the shaded area represents the 95% confidence intervals. The two sites are approximately 95 km apart, located as given in Materials and Methods. For comparison between sea surface temperatures offshore at each site see figure S4. **(B)** Comparison of the mean egg temperatures of the two species of turtle versus the average of sea surface temperatures taken between 15 June-15 July of each year.

**Figure 4.**
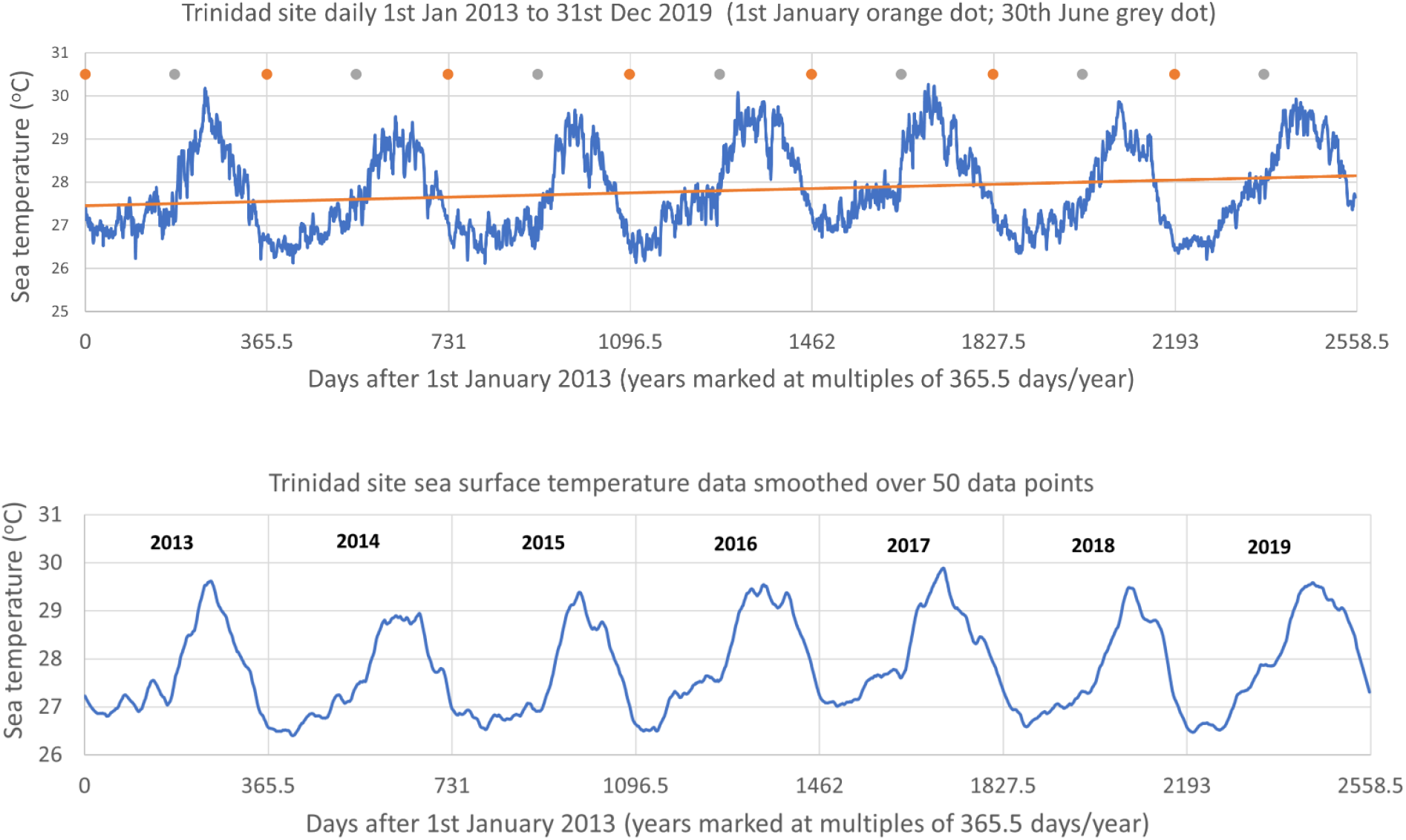
Seasonal sea surface temperature fluctuations offshore from the Trinidad leatherback turtle nest beach. The daily records for the Trinidad sites with dates as indicated for each year, and below, the same data smoothed over fifty data points. Some of the smaller peaks (clearest in the smoothed data), in 1913 for instance, appear to coincide with full Moons and therefore consequent tidal changes. Note the shoulders of warmer water appearing earlier before the summer peaks in 2016 and 2017 but not in the other years. The fitted line is a linear regression line fitted across all the data. See figure S4 for similar plots for the Tobago hawksbill turtle nesting site.

Our observation period encompassed one strong El Niño year (the warm phase of the El Niño–Southern Oscillation (ENSO)) between 2015 and 2016, two weaker El Niños, and two weak La Niña events (the cool phase of the ENSO), as indicated in figure 2. There is insufficient information to discern any long-term patterns at this stage, or connection between ENSO and waters local to our observation sites, but it is noticeable that, for hawksbills, the strong 2015-2016 El Niño event was accompanied by an increase in core temperatures, the two following weak La Niñas with a reduction, and the two subsequent weak El Niños with slight increases, over a previous year.

## 4. Discussion

We found notable year-to-year changes in the core body temperatures of two species of sea turtle with contrasting heat-conserving anatomies, physiologies, and migratory patterns, nesting on beaches in the same ocean area. If local sea and thence turtle core temperatures are pertinent to nest site selection as global ocean warming proceeds, then it will be important to understand such year-to-year fluctuations across different nesting areas. The use of non-contact infrared thermometers, as used here, is effective for large-scale surveys that could detect differences between beaches and trends on individual beaches.

A particular reason to consider body temperatures of turtles coming ashore is potentially to detect thermally deleterious challnges in the approaches to currently-used nesting beaches. Since we do not know the offshore waters frequented by turtles before coming ashore, their body temperatures could provide information on pertinent changes. For instance, whether offshore waters remain thermally tolerable may be moot for some Pacific Ocean populations of sea turtles given the increasingly globally influential periodic El Niño warming events, and unusually high temperature east-to-west bulk water flow across the equatorial Pacific. Whether water temperatures will become intolerable or not to some species remains controversial given current theoretical models that are perforce based on untested assumptions of critical body temperatures in sea turtles, and limitations to predictions of ocean warming. But changes in body temperatures of nesting turtles, readily obtained, as here, may provide clues as to future viability of nesting beaches.

Charting the sea surface temperatures adjacent to both of our nesting sites shows regular annual cycles within which there was a gradual overall increase during our study period (figures 4 and S4). Surprisingly, therefore, the surface temperatures of the turtle eggs showed an overall slight decrease during the study years (figure 2). This disparity could have been due to inconsistencies in how measurements were taken (though none such were apparent), or changes in the instruments used (though there were several IR thermometers recruited over the years, all calibrated yearly). The incidental timings of our nocturnal field measurements could also have an effect on the overall trend seen; lunar cycles and tidal effects are known to affect the frequency of turtles coming ashore [35] (lunar dates would, of course, change from year to year), and, as seen in figures 4 and S4, some peaks in temperatures appear to correspond to full Moons, tides presumably changing the temperature of near shore waters. Also, in the years 2016 and 2017 warmer waters begin to appear before the summer peak earlier than in other years (figures 4 and S4). Nevbertheless, the overall pattern of core temperature changes of both leatherbacks and hawksbills from year to year did approximately follow sea surface temperatures – the drops in June and July sea surface temperatures seen in 2015 and 2018 correspond to minima for both species of turtle core temperatures (figure 3B). Sampling inconsistencies aside, more interesting would be if changes in sea currents local to the study beaches were occurring, or the turtles were changing where they were resting, mating, or feeding before coming ashore. Also notable is that the core temperatures of the hawksbill turtles were always above that of the sea surface temperatures (figure 3B), meaning that this species retains some metabolic heat, or that they reside in waters higher in temperature than the sea surface from where we chose to collect satellite data, or that the temperature at the surface as measured by satellite is reduced by evaporation, wave action, wind, and so on, such that the water beneath is warmer.

Understanding how sea turtles may adapt their phenology of nesting in response to ocean warming is important, especially given the changes that have already been observed [14, 28, 34, 46]. Superficially, it would seem feasible for some species to exploit seasonal changes in to move to cooler times. This may apply to hawksbills, for instance, but leatherbacks may be more limited in choice given their presumed need to migrate north in winter for resources of abundant Cnidarians upon which to feed prior to breeding [47-51]. The other obvious change would be to move away from hot central tropical beaches. Sea turtles tend to return to their natal coastal areas to nest, but fidelity to beaches is not absolute [52, 53]. A degree of non-fidelity, behaviourally built-in or environmentally-induced, could be important to allow turtles to change nesting sites to improve their fitness. If the temperature of offshore waters were pertinent to female turtles’ selection of beaches to nest, then surveying core body temperatures of females could be predictive of whether a rookery will continue to be preferred. This, of course, would be separate from whether the beaches themselves remain good for egg development, hatchling emergence, and balanced temperature-dependent sex ratios [25, 27, 28].

On a longer timescale, it has been predicted that global warming will result in large land vertebrates becoming depleted in tropical areas [54], an effect from which the oceans will not be exempt [55]. Previous geologically brief, severe global warming events in the Eocene, for example, are deemed to have caused land mammals to have become smaller, allowing them to increase their shedding of heat from their surfaces [56]. And, on both land and sea, large vertebrates became depleted in the tropics during a time of increased global temperatures in the Triassic [57]. Moreover, during those Triassic warm periods the rate of reptile evolution accelerated [58], which could be encouraging for sea turtle species survival although the Triassic climatic changes would have been considerably slower than is currently the case. While the future scale, rate, and limits of global warming remains to be seen, it is conceivable that some tropical waters and nesting beaches will become too hot for breeding sea turtles to tolerate and their eggs to develop, such that successful rookeries will diverge poleward. Of pertinence, a recent survey has shown that many groups of animals are already undergoing poleward movements [59]. It may therefore be that discrete subpopulations of sea turtles develop to north and south of equatorial seas that will no longer interbreed. Given the differing scale and distribution of temperature anomalies in the Atlantic and Indo-Pacific oceans, such an effect may differ between them. If leatherback turtles were to evolve reduced body masses in order to lose heat more effectively in warming tropical seas, how would that then affect their survival in their temperate to near sub-polar winter feeding grounds?

For now, our finding of year-to-year fluctuations in body temperatures of nesting sea turtles using a simple method for large-scale surveys has potential implications for understanding their future environmental tolerances, migrations, nest site selection, phenology, and reproductive success. Similar long-term surveys for other rookeries may be useful to predict their long-term viability as nesting sites and how this will drive future changes in the survival and distribution of breeding sea turtles, perhaps even impinging on the evolution of their anatomies and physiologies.

## Ethics

All procedures and permissions to be on protected beaches and study nesting sea turtles were approved by the Wildlife Division of the Government of Trinidad and Tobago.

## Data accessibility

Supplementary A contains figures S1 to S4, and Tables S1A and S1B. Supplementary B contains the primary egg surface temperature data used for the graphs and statistical analyses. Supplementary C presents the satellite-based sea surface temperature data obtained as described in Methods. All supplements are available in the Dryad data repository at https://doi.org/10.5061/dryad.0gb5mkm6p.

## Authors’ contributions

MWK planned the study, supervised the observations, curated and analysed the data, administered the project, acquired resources, created the figures, and wrote the paper.

## Acknowledgements

We thank the Wildlife Section of the Trinidad and Tobago Government for access to nesting beaches, and the beach patrolling teams in Fishing Pond in Trinidad, and North East Sea Turtles (NEST), Tobago, for their field support and assistance. We are particularly grateful to Sookraj Persad in Trinidad, and Devon Eastman and Andel Mackenzie in Tobago, for passing on their considerable experience on turtle nesting and assisting with field observations. We also thank members of the separate Glasgow University expeditions to Trinidad and Tobago from 2013 to 2019 for their dedicated assistance in the field, and Isobel Byrne and Duncan Murray-Uren for provision of the 2019 Tobago hawksbill data. Particular thanks go to Dr Tomas J Burns with whom this study was originally conceived and initiated, and with whom the field data were obtained in the years 2013 to 2015.

