## Supplement A for "A seven-year record of fluctuating core body temperatures of nesting leatherback and hawksbill sea turtles"

Malcolm W. Kennedy

School of Biodiversity, One Health & Veterinary Medicine, College of Medical, Veterinary and Life Sciences, Joseph Black Building, University of Glasgow, Glasgow G12 8QQ, Scotland, UK.

**Supplementary A**

**Figure S1. Core body temperature differences between leatherback and hawksbill turtles.**

**Table S1. ANOVA comparison of year-to-year variation in core body temperatures of leatherback and hawksbill sea turtles.**

**Figure S2. Changes in turtle core body temperatures during each of the seven sampling seasons.**

**Figure S3. Comparison of sea temperatures between the Trinidad and Tobago sites.**

**Figure S4. Daily sea surface temperatures measured offshore of the Trinidad and Tobago sites from 2013 to 2019.**

**The original/raw data for all of the graphs and statistical analyses presented are available in Supplementaries B and C**

**Figure S1. Core body temperature differences between leatherback and hawksbill turtles.** Data point and Kennel smooth distribution companion (with Mean + SD indicated) to the box and whisker representation of the data presented in figure 1. Cumulative egg temperature measurements of the two species over the seven field seasons 2013 – 2019 inclusive. Data were obtained from 271 nesting instances of leatherback and 312 hawksbill turtles.

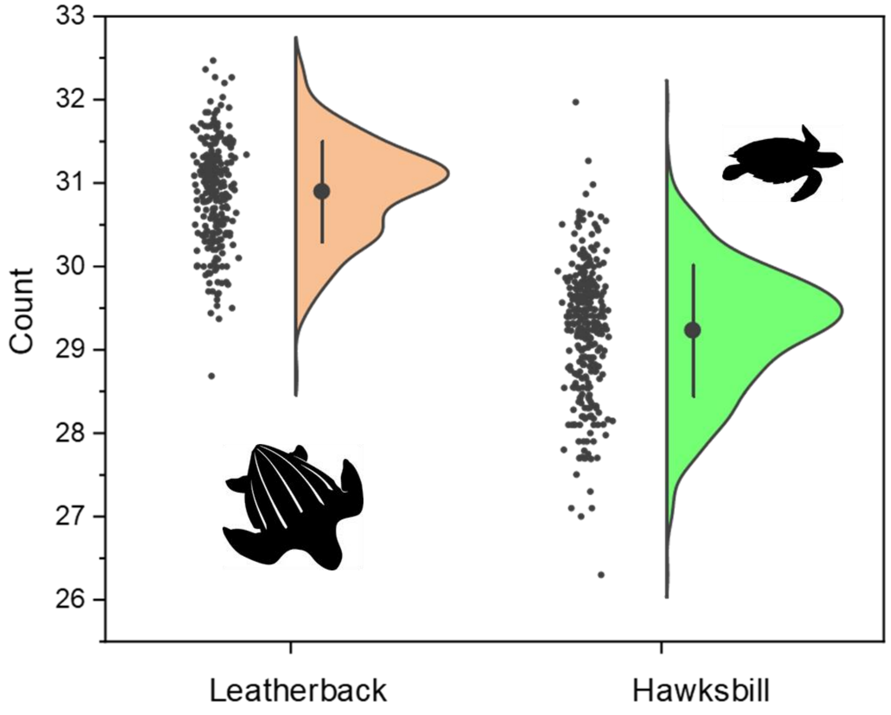

**Table S1. ANOVA comparison of year-to-year variation in core body temperatures of leatherback and hawksbill sea turtles.** Analysis of variance analysis with the Tukey procedure at 95% confidence. Grouping tables where means that do not share a letter are significantly different ($p < 0.05$).

**Leatherback turtles**

| Year | N | Mean | Grouping | | |
|---|---|---|---|---|---|
| 2013 | 43 | 31.49 | A | | |
| 2014 | 22 | 31.27 | A | B | |
| 2017 | 24 | 31.17 | A | B | |
| 2016 | 56 | 31.07 | | B | |
| 2015 | 27 | 30.98 | | B | |
| 2019 | 39 | 30.40 | | | C |
| 2018 | 60 | 30.37 | | | C |

**Hawksbill turtles**

| Factor | N | Mean | Grouping | | | | | |
|---|---|---|---|---|---|---|---|---|
| 2013 | 18 | 30.31 | A | | | | | |
| 2016 | 63 | 29.91 | A | B | | | | |
| 2014 | 11 | 29.43 | | B | C | D | | |
| 2015 | 82 | 29.31 | | | C | | | |
| 2017 | 90 | 28.99 | | | | D | E | |
| 2019 | 14 | 28.56 | | | | | E | F |
| 2018 | 34 | 28.10 | | | | | | F |

**Figure S2. Changes in turtle core body temperatures during each of the seven sampling seasons.** Measurements were taken in June, July and early August in the years indicated and the turtle numbers are given in the order they were sampled during the field season. Each point represents the average of the several recordings of individual freshly laid eggs made from each turtle. See Supplementary B for the complete data.

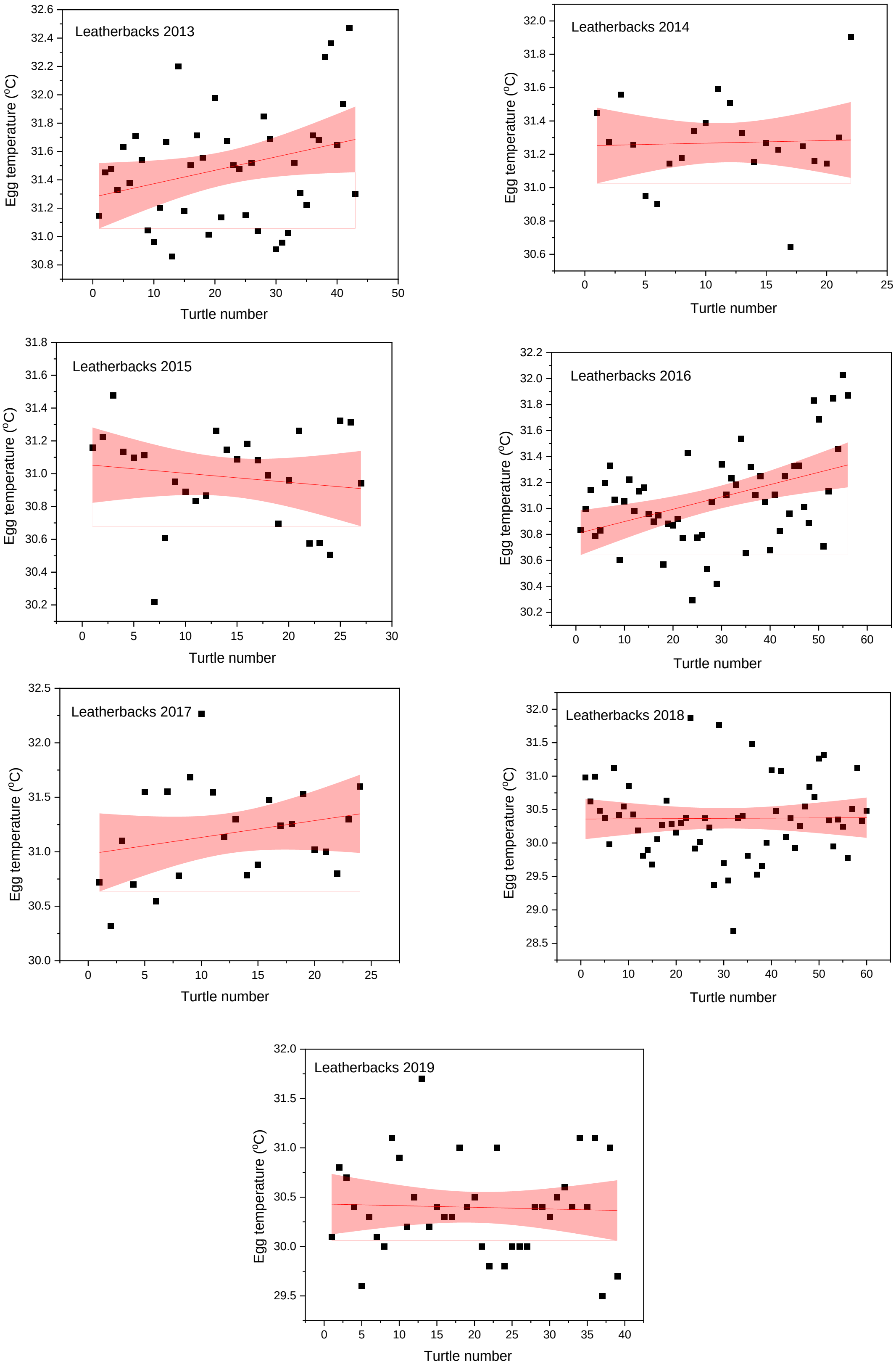

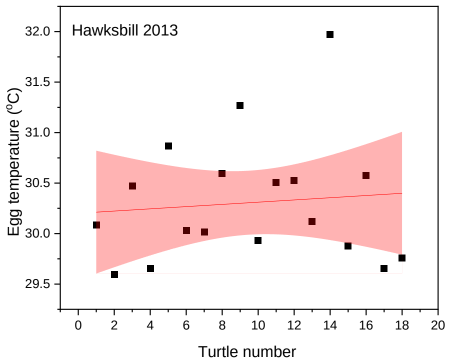
Hawksbill 2013
Egg temperature (ºC)
Turtle number

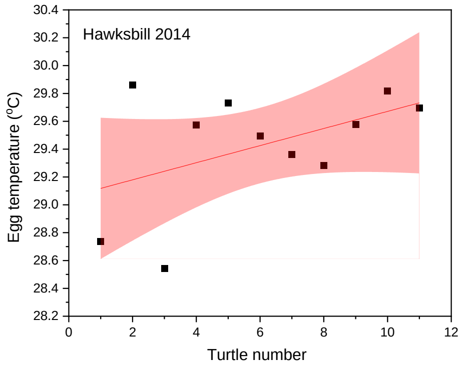
Hawksbill 2014
Egg temperature (ºC)
Turtle number

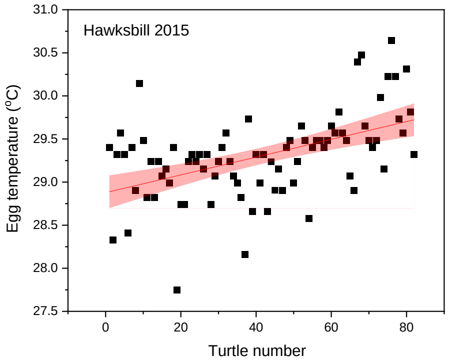
Hawksbill 2015
Egg temperature (ºC)
Turtle number

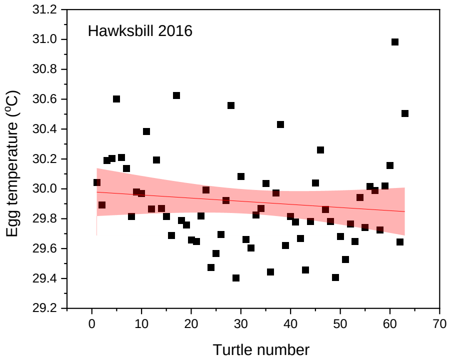
Hawksbill 2016
Egg temperature (ºC)
Turtle number

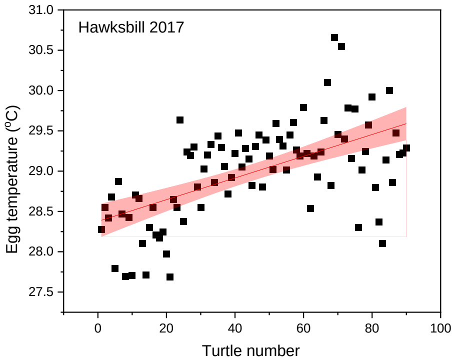
Hawksbill 2017
Egg temperature (ºC)
Turtle number

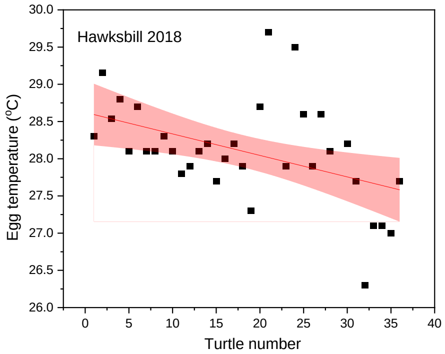
Hawksbill 2018
Egg temperature (ºC)
Turtle number

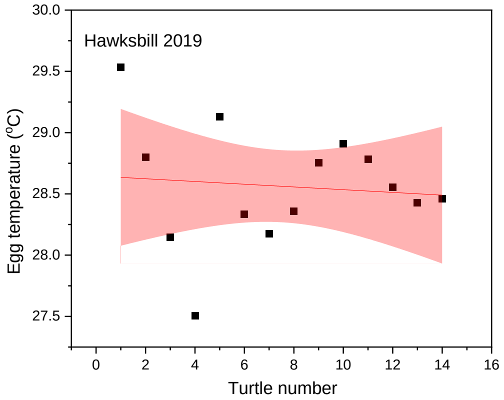
Hawksbill 2019
Egg temperature (ºC)
Turtle number

**Figure S3. Comparison of sea temperatures between the Trinidad and Tobago sites.** Data points are the averages of daily sea surface temperatures taken offshore at positions detailed in Materials and Methods between June 15th and July 15th of each year. Linear fitting yields slope $y = 0.7605x + 6.5669$; $R^2 = 0.9312$.

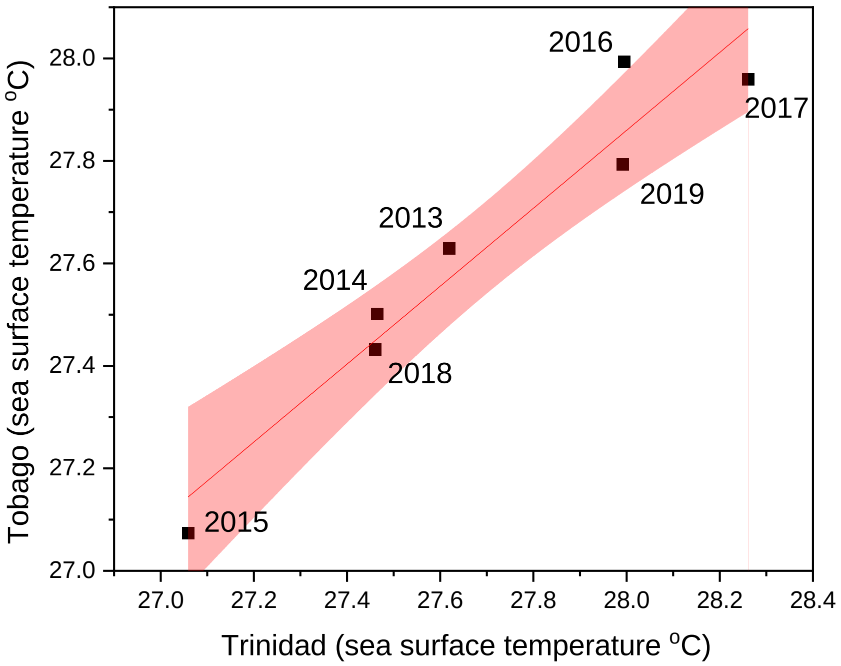

**Figure S4. Daily sea surface temperatures measured offshore of the Trinidad and Tobago sites from 2013 to 2019.** Sea surface temperatures were obtained from satellite data centered offshore as described in Mathods. **(A)** The daily records for the Trinidad sites with dates as indicated for each year. **(B)** The same data as in (A) but smoothed over fifty data points using default smoothing procedure in ORIGINlab software. **(C)** and **(D)**, as for (A) and (B) above but for the Tobago site. Some of the smaller peaks (clearest in the smoothed data), in 1913 for instance, appear to coincide with full Moons and therefore consequent tidal changes. Note also the shoulders of warmer water appearing at both sites in 2016 and 2017 in the period before June 30$^{th}$ but not to nearly the same extent in the other years.

**A**

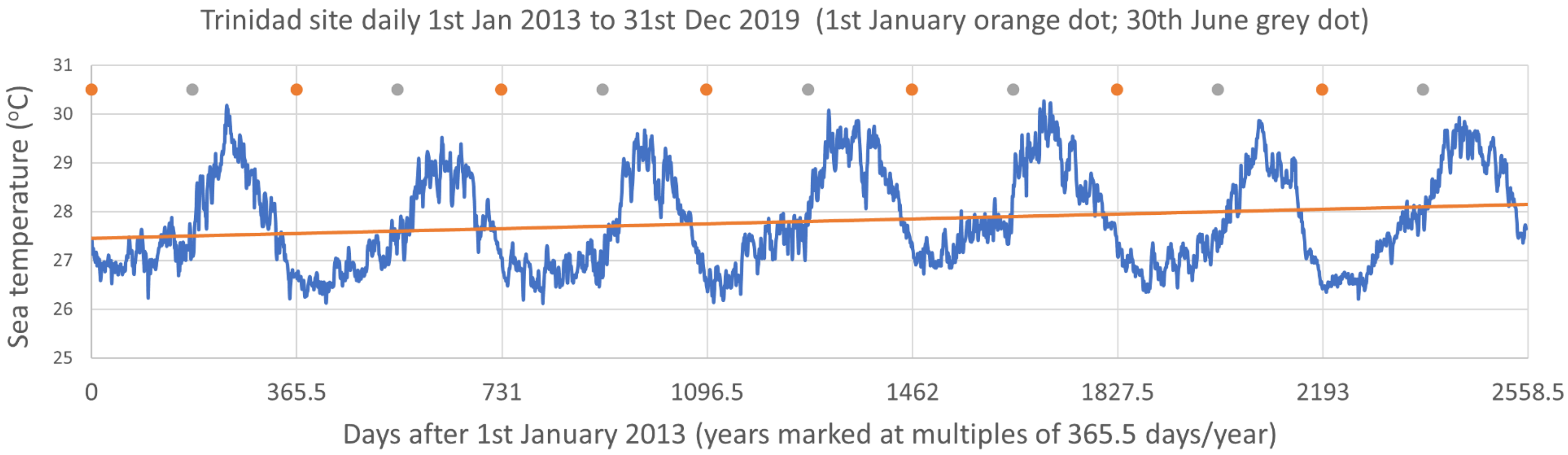
Trinidad site daily 1st Jan 2013 to 31st Dec 2019 (1st January orange dot; 30th June grey dot)
Sea temperature (°C)
31
30
29
28
27
26
25
0
365.5
731
1096.5
1462
1827.5
2193
2558.5
Days after 1st January 2013 (years marked at multiples of 365.5 days/year)

**B**

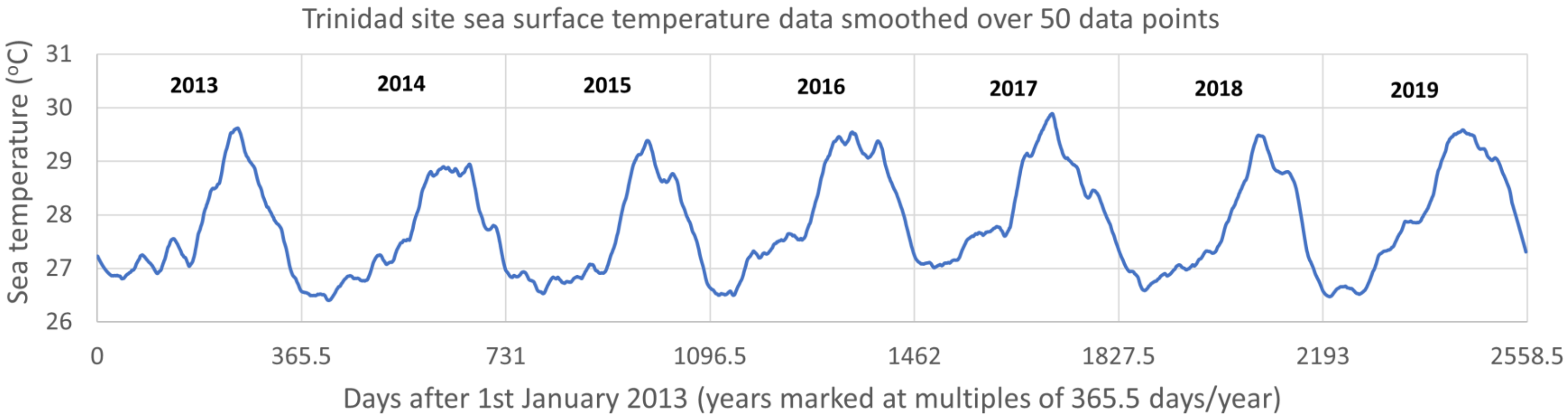
Trinidad site sea surface temperature data smoothed over 50 data points
Sea temperature (°C)
31
30
29
28
27
26
2013
2014
2015
2016
2017
2018
2019
0
365.5
731
1096.5
1462
1827.5
2193
2558.5
Days after 1st January 2013 (years marked at multiples of 365.5 days/year)

**C**

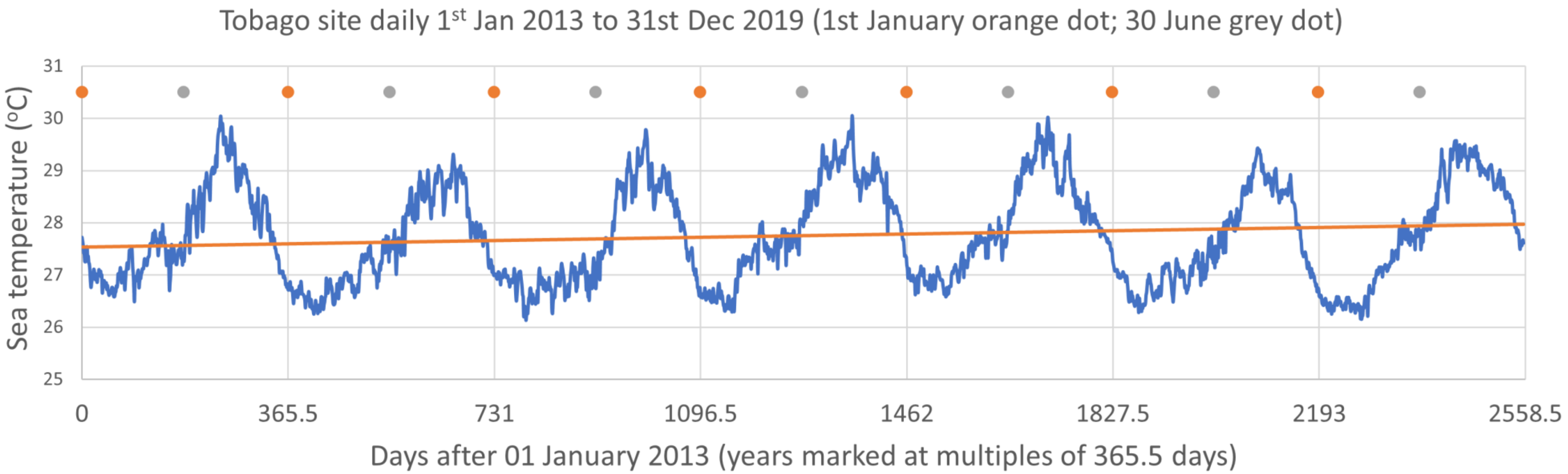

Tobago site daily 1st Jan 2013 to 31st Dec 2019 (1st January orange dot; 30 June grey dot)
Sea temperature (°C)
31
30
29
28
27
26
25
0
365.5
731
1096.5
1462
1827.5
2193
2558.5
Days after 01 January 2013 (years marked at multiples of 365.5 days)


**D**

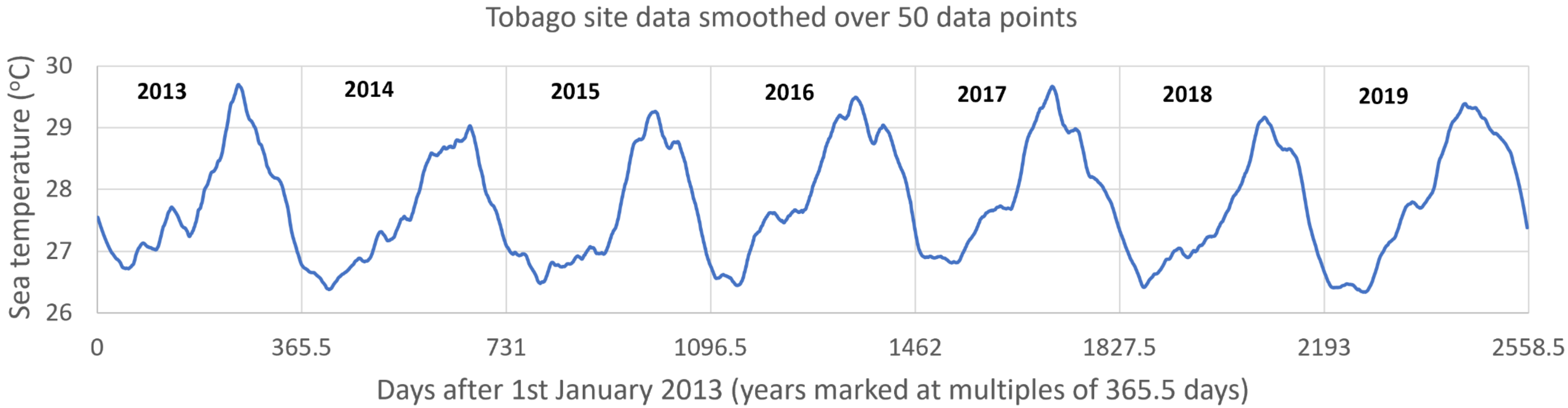

Tobago site data smoothed over 50 data points
Sea temperature (°C)
30
29
28
27
26
2013
2014
2015
2016
2017
2018
2019
0
365.5
731
1096.5
1462
1827.5
2193
2558.5
Days after 1st January 2013 (years marked at multiples of 365.5 days)
